# A pan-cancer metabolic atlas of the tumor microenvironment

**DOI:** 10.1101/2020.10.16.342519

**Authors:** Neha Rohatgi, Umesh Ghoshdastider, Probhonjon Baruah, Anders Jacobsen Skanderup

## Abstract

Tumors are heterogeneous cellular environments with entwined metabolic dependencies. Here, we used a tumor transcriptome deconvolution approach to profile the metabolic states of cancer and non-cancer (stromal) cells in bulk tumors of 20 solid tumor types. We identified metabolic genes and processes recurrently altered in cancer cells across tumor types, including pan-cancer upregulation of deoxythymidine triphosphate (dTTP) production. In contrast, the tryptophan catabolism rate limiting enzymes, *IDO1* and *TDO2*, were highly overexpressed in stroma, suggesting that kynurenine-mediated suppression of antitumor immunity is predominantly constrained by the stroma. Oxidative phosphorylation was unexpectedly the most upregulated metabolic process in cancer cells compared to both stromal cells and a large atlas of cancer cell lines, suggesting that the Warburg effect may be less pronounced in cancer cells *in vivo*. Overall, our analysis highlights fundamental differences in metabolic states of cancer and stromal cells inside tumors and establishes a pan-cancer resource to interrogate tumor metabolism.

## Introduction

Malignant cells reprogram metabolic processes to support their uncontrolled growth and proliferation, which may be exploited to diagnose and treat tumors (Vander Heiden and DeBerardinis 2017). The tumor microenvironment (TME) contains extracellular matrix, myofibroblasts, immune cells and vascular networks, collectively referred to as stroma (Chen et al. 2015). Cancer cells can acquire nutrients and growth supporting metabolites from the stroma (Sousa et al. 2016). Additionally, secreted metabolites from cancer cells can serve as paracrine factors to direct the fate of stromal cells (Schwörer, Vardhana, and Thompson 2019). However, our understanding of cancer and stromal cell metabolism inside tumors and their joint influence on tumor growth and progression is still nascent (Hyduke, Lewis, and Palsson 2013; Resendis-Antonio et al. 2015).

Metabolic alterations have been widely accepted as a hallmark of cancer, however systematic large-scale profiling of tumor metabolic phenotypes remains uncommon due to technical challenges in identifying and measuring metabolites (Reznik et al. 2018; Hanahan and Weinberg 2011). Global metabolomic profiling of tumor tissues have linked metabolites such as 2-hydroxyglutarate, cysteine, kynurenine, lipids, and fatty acids with tumor progression in distinct tumor types (Hakimi et al. 2016; Kamphorst et al. 2015; Zhang et al. 2013; Budhu et al. 2013; Huang et al. 2013; Prabhu et al. 2014). Meta-analyses have shown that metabolites such as lactate, kynurenine and taurine, are repeatedly upregulated in tumors as compared to normal tissue across tumors of different origin (Reznik et al. 2018; Goveia et al. 2016). Gene expression data has also been used to interrogate the metabolic phenotypes of tumor tissue (Nilsson et al. 2014; Hu et al. 2013; Hakimi et al. 2016). Single cell gene expression analysis of inferred malignant and non-malignant cells can additionally be used to study intra-tumor heterogeneity of metabolic pathways (Xiao, Dai, and Locasale 2019). Finally, systematic *in vitro* metabolomic profiling across a large compendium of cancer cell lines has linked metabolic pathway activities to genetic and epigenetic alterations in cultured cancer cells (Li et al. 2019).

Despite these existing large-scale studies of tumors, we still only have a limited description of how the metabolic programs and dependencies of cancer cells differ from their neighboring stromal cells inside the TME. Similarly, it is not clear to what extent the metabolic programs of cancer cells might change as they are removed from their natural milieu and cultured in the lab. To address these questions, we used a data-driven tumor transcriptome deconvolution approach (Ghoshdastider et al. 2019) to profile and compare cancer and stromal cell metabolic pathways across 7865 tumors and 20 solid tumor types. Using this approach, we explored the distinct metabolic signatures of cancer and stromal cells conserved across most solid tumor types. Additionally, using a large compendium of cancer cell line transcriptomes, we contrasted *in vivo* and *in vitro* cancer cell transcriptomes to analyze how cancer cells might adapt and change their metabolic programming in cell culture. All data generated in this study are available for interactive exploration and analysis at https://camp.genome.sg/.

## Results

### Overview of approach

We first obtained transcriptome data for 7865 tumor and 815 normal tissue samples from The Cancer Genome Atlas (TCGA) and the Genotype-Tissue Expression (GTEX) database, respectively (Fig. 1, Suppl. Table 1). A genome-scale model for human metabolism, RECON3 (Brunk et al. 2018), was used to filter and reduce the dataset to 1845 metabolic genes. We used a tumor transcriptome deconvolution approach to estimate expression of metabolic genes in cancer and stromal (non-cancer) cells (Ghoshdastider et al. 2019). Briefly, we estimated tumor purity (cancer cell fraction) using genomic and transcriptomic data from each tumor sample, followed by non-negative linear regression to infer the expression of individual metabolic genes in cancer and stromal cells of each tumor type (Fig. 1, Methods). The metabolic phenotypes of cancer and stromal cells were then compared and contrasted to matched normal tissues samples. Additionally, cancer cell expression *in vivo* was compared to expression profiles of 620 cancer cell lines obtained from the cancer cell line encyclopedia (CCLE).

**Figure 1:**
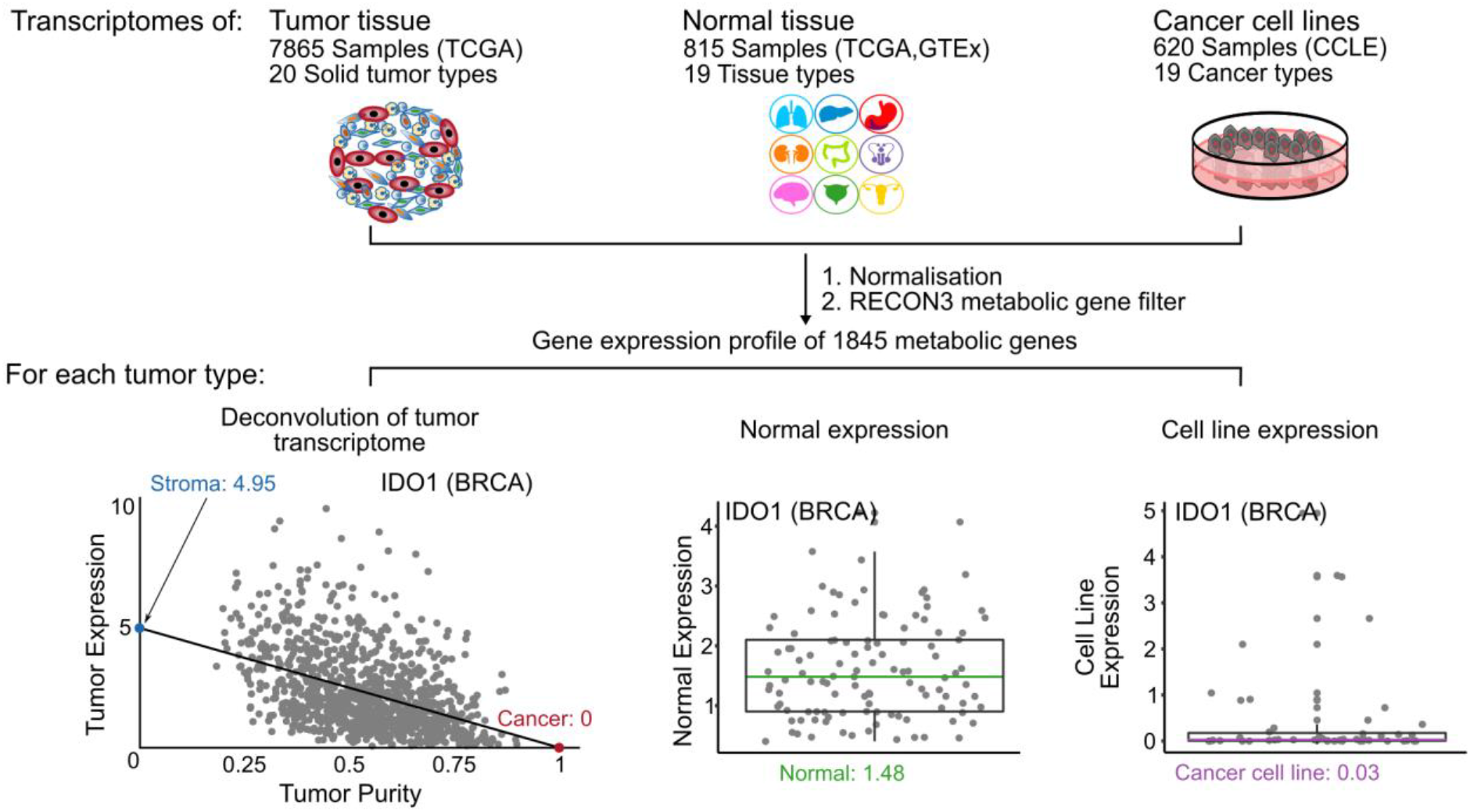
Overview of approach. Aggregation and comparison of transcriptome metabolic profiles across cancer cells, stromal cells, normal tissue and cancer cell lines across different tissue types. For each tumor type, cancer and stromal cell gene expression was inferred using a tumor transcriptome deconvolution approach, regressing bulk tumor expression against estimated tumor purity.

### Discovery of cell-type specific metabolic gene expression

To gain insights into conserved metabolic phenotypes of cancer and stromal cells across tumor types, we compared the expression levels of individual metabolic genes across cancer cells, stromal cells, and normal tissue. We first calculated the cell type specificity for each gene as the ratio of expression in a given cell type divided by the total expression for all three cell types (i.e., cancer, stroma and normal cells, Methods). For each gene, we then computed the median cell type specificity across tumor types. This analysis revealed a limited number of metabolic genes with high cancer cell specificity across tumor types (28 genes with >60% cancer relative expression, Fig. 2a, b). In contrast, we observed a large number of genes with high stromal specificity across most tumor types (93 genes with >60% stromal relative expression, Fig. 2a). This observation possibly reflects that cancer cell metabolism is governed by tissue and cell lineage, whereas tumor-infiltrating stromal cells comprise a more conserved collection of cell types.

**Figure 2:**
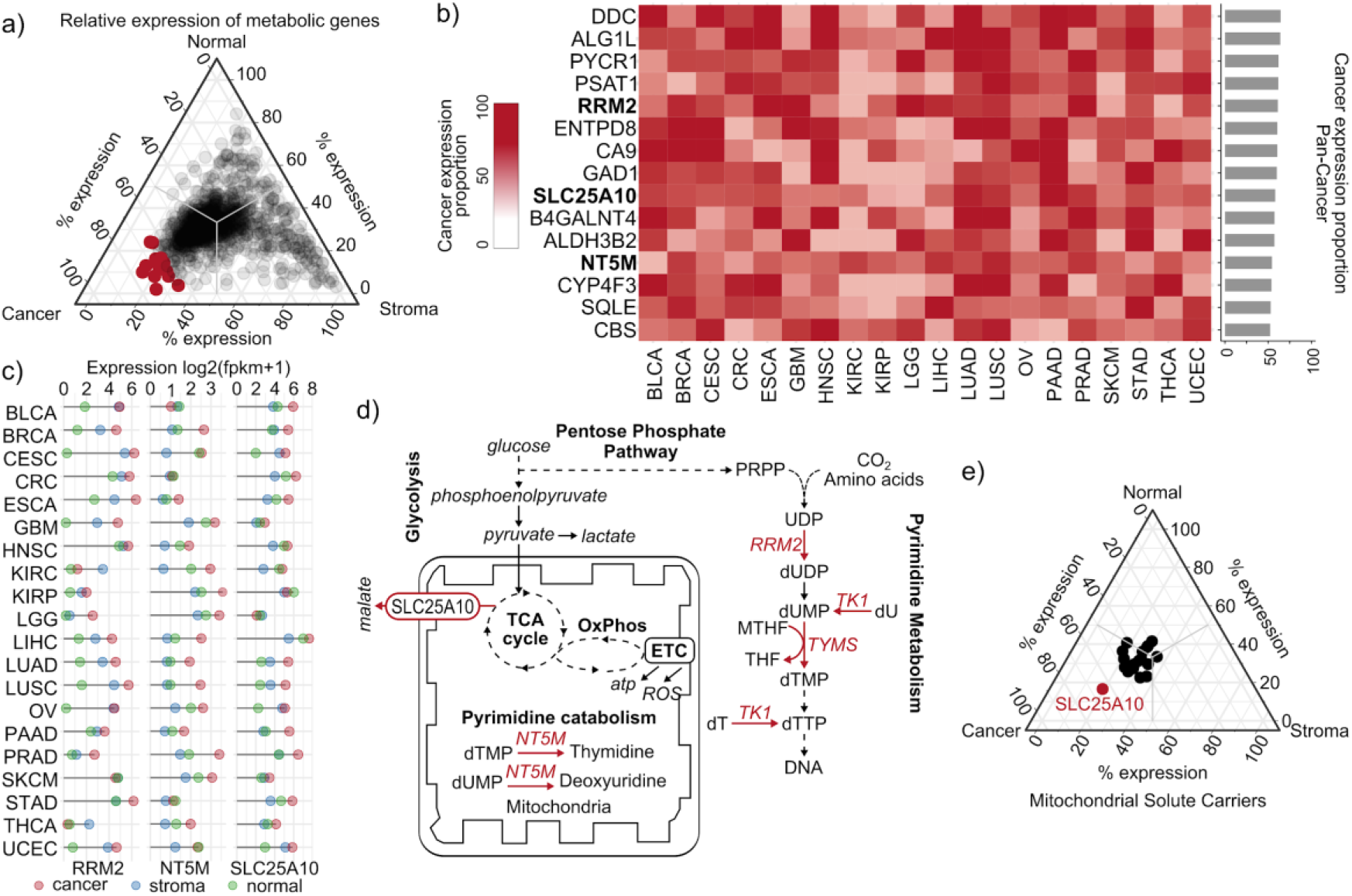
Cancer cell specific expression of metabolic genes across tumor types. **a**) Relative expression (proportion) of metabolic genes in cancer, stroma and corresponding normal tissue across tumor types. The top-15 genes overexpressed in cancer cells are highlighted (red). **b**) Relative expression profiles of the top-15 overexpressed genes in cancer cells. **c**) Expression of *RRM2*, *NT5M* and *SLC2A10* in cancer, stromal and normal cells across tumor types. **d**) Metabolic pathway map highlighting cancer cell upregulation of dTTP metabolism and *SLC25A10*, transporting dicarboxylates across the mitochondrial membrane. **e)** Relative gene expression of mitochondrial solute carrier proteins in cancer, stroma and normal cells.

### Pyrimidine synthesis capacity is upregulated in cancer cells

We next explored the functional characteristics of cancer-cell specific metabolic genes. Gene ontology enrichment analysis of genes with >60% cancer cell-specificity (N=28), showed enrichment of genes related to DNA metabolism (Fisher-exact test, 5/27 genes, adj. *P* = 0.04, Suppl. Table 2). Strikingly, 3 genes in this pathway (*RRM2*, *TK1* and *TYMS*) are involved in pyrimidine synthesis. The regulatory subunit of ribonucleotide reductase, *RRM2*, had one of the highest cancer cell gene expression specificities across tumor types (Fig. 2b). Ribonucleotide reductases catalyzes the rate-limiting step of de novo synthesis of all four deoxyribonucleotides (Mathews 2006). Our data suggests that *RRM2* has more than 8-fold higher expression in cancer cells as compared to matched normal tissue across 9 tumor types and more than 2-fold higher expression in 16 tumor types (Fig. 2b and 2c). *TYMS* (Thymidylate synthase) and *TK1* (thymidine kinase) catalyze the de novo and salvage synthesis of deoxythymidine triphosphate (dTTP) (Fig. 2d) in the nucleus at sites of DNA synthesis (Stover and Weiss 2012). We observed both *TYMS* and *TK1* have more than 4-fold higher expression in the cancer cells of 12 out of 20 tumor types as compared to the normal tissue. We next investigated if DNA copy number amplifications could be driving upregulation of *RRM2, TK1* and *TYMS* expression. We found that these genes were affected by both DNA copy number gain and loss across tumors of different tumor types (Suppl. Fig. 1 & 2). However, the inferred cancer cell expression was consistently higher than stromal and normal cells irrespective of DNA copy number alterations, suggesting that the upregulation of these genes may instead be mediated by epigenetic or transcriptional regulation. Overall, this suggests that cancer cells have increased capacity for dTTP production.

### Dysregulation of mitochondrial metabolic pathways in cancer cells

Dysregulation of metabolic processes in mitochondria may contribute to tumorigenesis (Neagu et al. 2019; Porporato et al. 2018). Interestingly, among the genes with highest cancer cell expression specificity across tumor types, we identified the mitochondrial 5’ nucleotidase (*NT5M*). In 15/20 tumor types, *NT5M* had >2 fold increase in cancer cell expression as compared to stromal cells (Fig 2b,c). Interestingly, the mitochondrial 5’ nucleotidase catalyzes the dephosphorylation of nucleoside triphosphates, especially dUMPs and dTMPs (Rampazzo et al. 2000). Since dTTP accumulation has been shown to result in defects in the mitochondrial genome (Rampazzo et al. 2000), it is conceivable that *NT5M* plays a protective role by maintaining mitochondrial dTTP homeostasis in the pyrimidine synthesizing cancer cells.

We also identified the mitochondrial dicarboxylate carrier *SLC25A10* (Fig. 2a). In 13/20 tumor types, this gene had >2 fold increase in cancer expression as compared to stromal cells (Fig. 2b). Pancreatic adenocarcinoma (PAAD) showed highest relative expression of SLC25A10, with >6-fold upregulation in cancer vs stromal cells (Fig. 2c). Analysis of all known mitochondrial transporters revealed that *SLC25A10* was the only transporter with significant pan-cancer upregulation in cancer cells (Fig. 2e), suggesting that *SLC25A10* upregulation is specific and not due to general upregulation of mitochondrial transporters in cancer cells. *SLC25A10* assists in the transport of dicarboxylates, such as malate and succinate across the mitochondrial membrane, thus supplying substrates for diverse processes such as gluconeogenesis, urea cycle and fatty acid synthesis (Fig. 2d). On investigating DNA copy number amplifications of *SLC25A10* and *NT5M,* we found that the inferred cancer cell expression was not affected by the DNA copy number alterations (Suppl. Fig. 1 & 2). Overall, this data suggests that cancer cells may promote mitochondrial transport to fuel specific anabolic processes like gluconeogenesis and fatty acid synthesis.

### Upregulation of IDO1 and kynurenine pathway in stroma across tumor types

Next, we investigated metabolic genes with stromal cell specific expression across tumor types. Indoleamine 2,3-dioxygenase 1 (*IDO1*) was among the genes most highly overexpressed in stromal cells across all tumor types (Fig. 3b,c, Suppl. Fig. 3 & 4). *IDO1* catalyzes the rate-limiting step in the kynurenine pathway converting tryptophan into N-formylkynurenine. Since depletion of tryptophan and accumulation kynurenine in the TME impairs T-cell functions (Xue et al. 2018; Fallarino et al. 2003; de Souza Sales et al. 2011), *IDO1* is being targeted in a number of clinical trials as an adjuvant with other cancer treatments (Zhai et al. 2018). Absolute stromal *IDO1* expression was highest in two gynecologic cancers (CESC and UCEC) and melanoma (SKCM) (Fig. 3c). The *IDO1* expression in the stromal cells of melanoma tumors is 7.01 Log2 (FPKM), whereas the expression in cancer cells is zero. Both the gynecologic cancers show a >8-fold upregulation of *IDO1* expression in stromal cells as compared to the cancer cells within the tumors. Strikingly, 19/20 tumor types show a more than 2-fold upregulation of *IDO1* expression in stromal cells as compared to the cancer cells. In cancer cells, *IDO1* expression was generally downregulated as compared to matched normal tissue (Fig. 3c, Suppl. Fig 4). 9 out of 20 tumor types show a more than 2-fold downregulation of *IDO1* expression in cancer cells as compared to the adjacent normal. Strikingly, further exploration of expression signatures along the tryptophan metabolism pathway, revealed frequent upregulation of *TDO2* and *IL4I1* in stroma (Fig. 3b). *TDO2* catalyzes the same reaction as *IDO1* in the kynurenine pathway and has a higher affinity for tryptophan as compared to *IDO1* (Ye et al. 2019). Interestingly, stromal cells showed >4-fold expression of *TDO2* in 12/20 and 8/20 tumor types as compared to cancer and normal cells, respectively. Other genes in the tryptophan metabolism pathway, *IL4I1*, *KYNU*, *KMO* and *CYP1B1* also showed >2-fold upregulation in stroma as compared to cancer cells in at least 18 out of 20 tumor types (Fig 3e). Together, these genes catalyze conversion of tryptophan to kynurenine.

**Figure 3:**
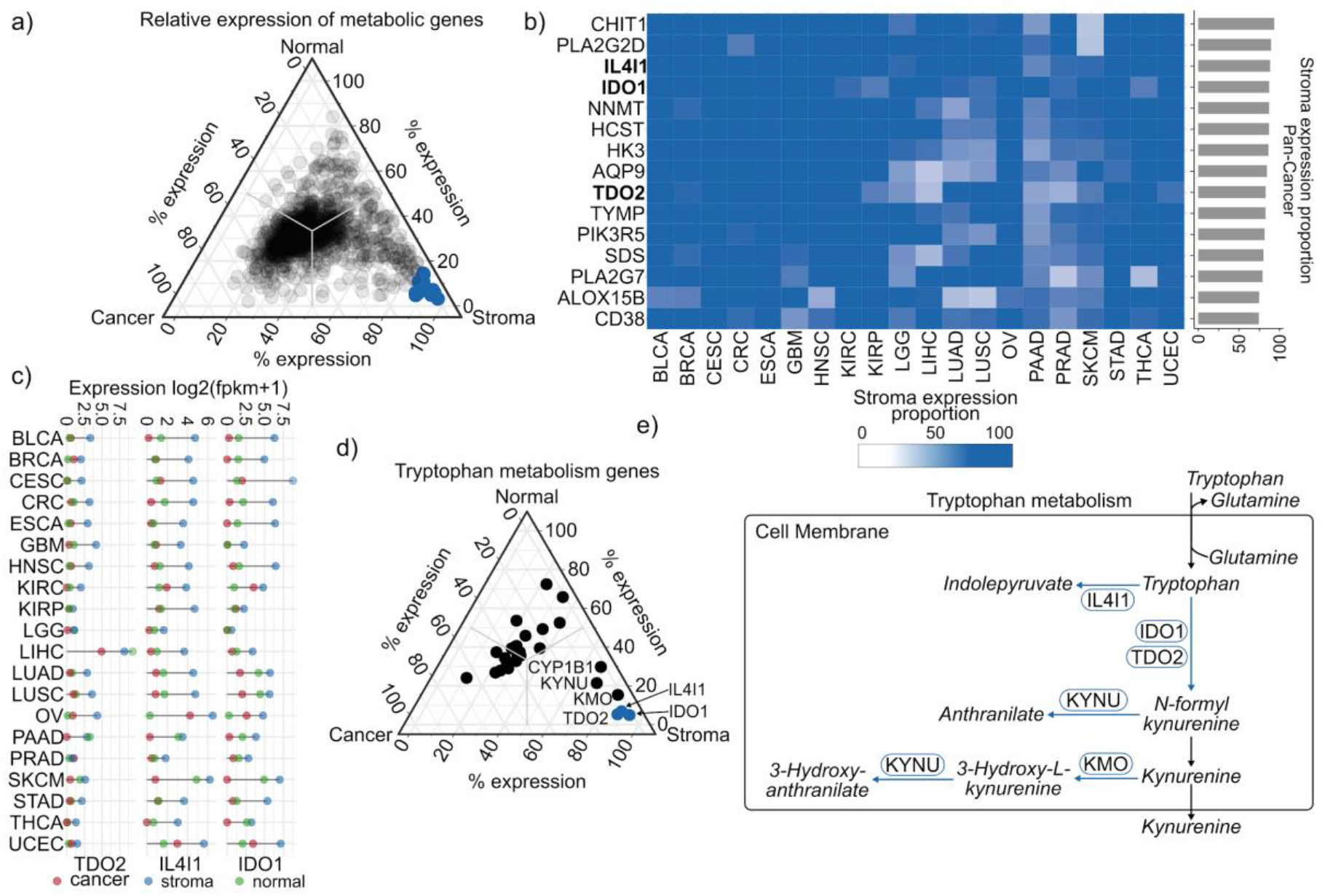
Stromal cell-type specificity of metabolic gene expression across tumor types. **a**) Relative expression (proportion) of metabolic genes in cancer, stroma and corresponding normal tissue across tumor types. The top-15 overexpressed stromal genes are highlighted in blue color. **b**) Relative expression profiles of the top overexpressed metabolic genes in stroma. **c**) Expression of *IDO1*, *TDO2* and *IL4I1* in cancer, stromal and normal cells across tumor types. **d)** Relative expression of metabolic genes involved in tryptophan metabolism. **e)** Metabolic map of tryptophan-kynurenine metabolism pathway, highlighting significantly differentially expressed genes in cancer and stromal cells.

### Cancer cells in tumors exhibit increased capacity for oxidative phosphorylation

To elucidate differences in metabolic pathways of cancer and stromal cells we performed gene set enrichment analysis of 186 KEGG pathways (see Methods). KEGG pathway IDs were used to filter out the metabolic pathways for our analysis. Confirming our previous observation (Fig. 3e), tryptophan metabolism was significantly upregulated in stromal cells as compared to cancer cells (FDR Adj. P = 0.02, GSEA permutation test) (Fig. 4a). Unexpectedly, the tricarboxylic acid cycle (TCA) and oxidative phosphorylation (Oxphos) pathways were highly upregulated in cancer cells across tumor types (Fig. 4a, Suppl. Fig. 5). Oxphos was significantly upregulated (FDR Adj. P = 0.0009, GSEA permutation test) in cancer cells of 13/20 tumor types, and 17/20 tumor types showed relative upregulation of Oxphos genes (Fig. 4b). Additionally, Oxphos was upregulated in cancer cells as compared to adjacent normal tissue in 14/20 tumor types (Fig. 4b). Overall, this data suggests that cancer cells inside tumors display an increased capacity for oxphos as compared to both stromal and normal tissue, which is in agreement with a newer paradigm stating that oxphos is required for tumor growth in many tumor types (DeBerardinis and Chandel 2016).

**Figure 4:**
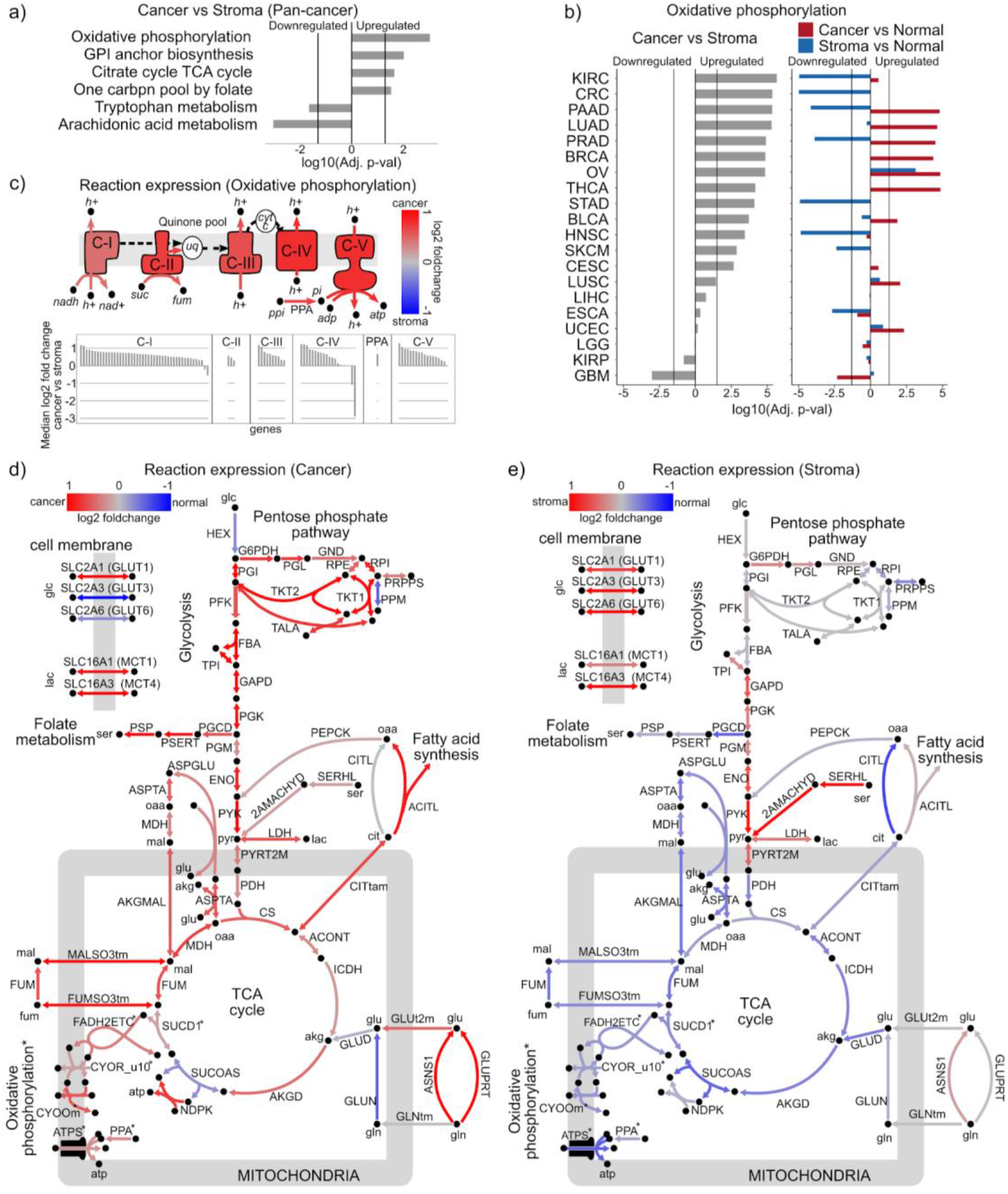
Differential metabolic pathway activity in cancer cells and stromal cells: **a)** Comparison of the KEGG metabolic gene sets in cancer vs stromal cells across tumor types. Out of 186 KEGG gene sets, 6 metabolic gene sets show significant difference after Benjamini-Hochberg FDR correction (Adj. *P* < 0.05). **b)** Enrichment of the KEGG oxphos pathway in cancer cells as compared to stromal cells across individual tumor types. **c)** Cancer vs. stroma activity estimated for individual reactions in the oxphos pathway. **d,e)** Metabolic maps showing reaction expression estimates of central carbon metabolism for cancer (**d**) and stromal (**e**) cells as compared to normal cells, across tumor types (excluding LGG and GBM).

Metabolic reactions are catalyzed by proteins that can be encoded by more than one gene. Thus, to further study Oxphos at the reaction level, we integrated deconvoluted cancer and stromal cell expression with a genome scale metabolic model, RECON3, using Gene-Protein-Reaction (GPR) associations present in this model (see Methods). GPRs use boolean annotation to describe genes encoding for protein complexes or isozymes that catalyse a particular reaction (Marín de Mas et al. 2019). Using the cancer and stroma specific metabolic models, we observed the differences in oxphos activity in the two compartments (Fig. 4c). Oxphos is performed by five complexes (Complexes I-V), first four of which are multi-subunit enzymes that create a protein gradient across the inner mitochondrial membrane that is used by complex V (ATP synthase) to generate ATP (Chaban, Boekema, and Dudkina 2014). Brain tumors (LGG and GBM) were excluded from this analysis because they showed a distinct trend from all other tumor types (Fig 4b & Suppl. Fig. 6). This analysis showed that all four electron transport complexes along with the ATP synthase were upregulated in cancer cells as compared to stromal cells (Fig. 4c). Analysis of the genes encoding the subunits of each complex confirmed increased metabolic capacity across all five Oxphos protein complexes in cancer cells as compared to stromal cells across the 18 tumor types (Fig. 4c).

### Stromal cells demonstrate Warburg-like phenotype

To further explore differences in central carbon metabolism (CCM) between cancer and stromal cells, we mapped cancer and stromal gene expression (relative to matched normal tissue) onto CCM metabolic maps generated from RECON3. This analysis highlighted an overall elevated metabolic capacity in CCM of cancer cells across different pathways: Glycolysis, Pentose phosphate pathway (PPP), TCA cycle and oxidative phosphorylation (Fig. 4d). *SLC2A1* (*GLUT1*), a membrane protein that transports glucose in almost all tissues under normal conditions, has previously been shown to be upregulated in many tumor types (Barron et al. 2016). Consistently, we observed >3 fold upregulation of *SLC2A1* in cancer cells and ~1.7 fold upregulation in stromal cells as compared to normal tissue (Fig. 4d & e). In contrast, *SLC2A3* (*GLUT3*) and *SLC2A6* (*GLUT6*), two other glucose transporters also with reported upregulation in a range of tumor types (Byrne et al. 2018; Barron et al. 2016), were downregulated in cancer cells by ~2.8 and ~1.2-fold, respectively, as compared to normal tissue (Fig. 4d). However, these two transporters were upregulated in stroma by ~2.6 and ~2.4-fold as compared to the normal tissue (Fig. 4e). Similarly, stromal cells of some tumor types (LIHC, KIRC, KIRP) also displayed >2-fold upregulation of hexokinase (HEX) reaction capacity, an enzyme that prepares glucose for glycolysis. These data indicate that, inside a tumor, stromal cells may have comparable, or even elevated, capacity to take up and process glucose as compared to cancer cells.

Next, we observed that the lactate dehydrogenase (LDH) reaction showed ~1.5 and ~1.3-fold upregulation in cancer and stromal cells, respectively, as compared to normal tissue, respectively. Additionally, lactate transporters *MCT1 (SLC16A1)* and *MCT4 (SLC16A3)* also showed upregulation in both cancer and stromal cells as compared to normal tissue across tumor types. The upregulation of *MCT1,* which is particularly suited for lactate uptake (Petersen et al. 2017), was comparable for cancer and stromal cells (1.3 vs. 1.4 fold). However, the expression of *MCT4*, the transporter more suited for lactate secretion, showed ~3.4 fold upregulation in stromal cells, with just ~1.8 fold upregulation in cancer cells (Fig. 4d & e). These observations further indicate that stromal cells, as compared to the cancer cells, could have a higher capacity to secrete lactate within the TME.

We also observed that reactions in the Malate aspartate shuttle (MAS) system, including *AKGMAL*, *MDH*, *ASPTA*, and *ASPGLU*, showed > 2 fold upregulation in cancer cells as compared to stromal cells of more than 5 out of 20 tumor types (Fig. 4d & 4e). The MAS utilizes exchangers to shuttle malate, alpha-ketoglutarate, oxaloacetate and aspartate in and out of the inner mitochondrial membrane in order to transfer NADH from cytosol to the mitochondrial matrix, which can be used to generate ATP in the electron transport chain. In combination with the elevated capacity of the TCA cycle in cancer cells (Fig 4d), this further highlights that cancer cells, in contrast to stromal cells, exhibit higher capacity for energy generation through oxidative phosphorylation and the electron transport chain. Overall, these results highlight that stromal cells have a high activity of glucose oxidation via glycolysis and lower activity of oxphos and thus, are in agreement with reverse Warburg theory (Pavlides et al. 2009; Fu et al. 2017; Martinez-Outschoorn et al. 2017).

### Cancer cells *in vitro* demonstrate downregulation of oxidative phosphorylation

To further explore metabolic differences between cancer cells *in vivo* and *in vitro*, we compared the deconvoluted cancer cell expression estimates in tumors with corresponding cancer cell lines from the Cancer Cell Line Encyclopedia (CCLE) (Li et al. 2019). We first calculated the expression specificity for each gene as the ratio of expression in a given cell type divided by the total expression for all three cell types (cancer *in vitro*, cancer *in vivo*, and normal tissue, Methods). This analysis revealed several genes with high *in vitro* gene expression specificity (>60%, Fig 5a).

**Figure 5:**
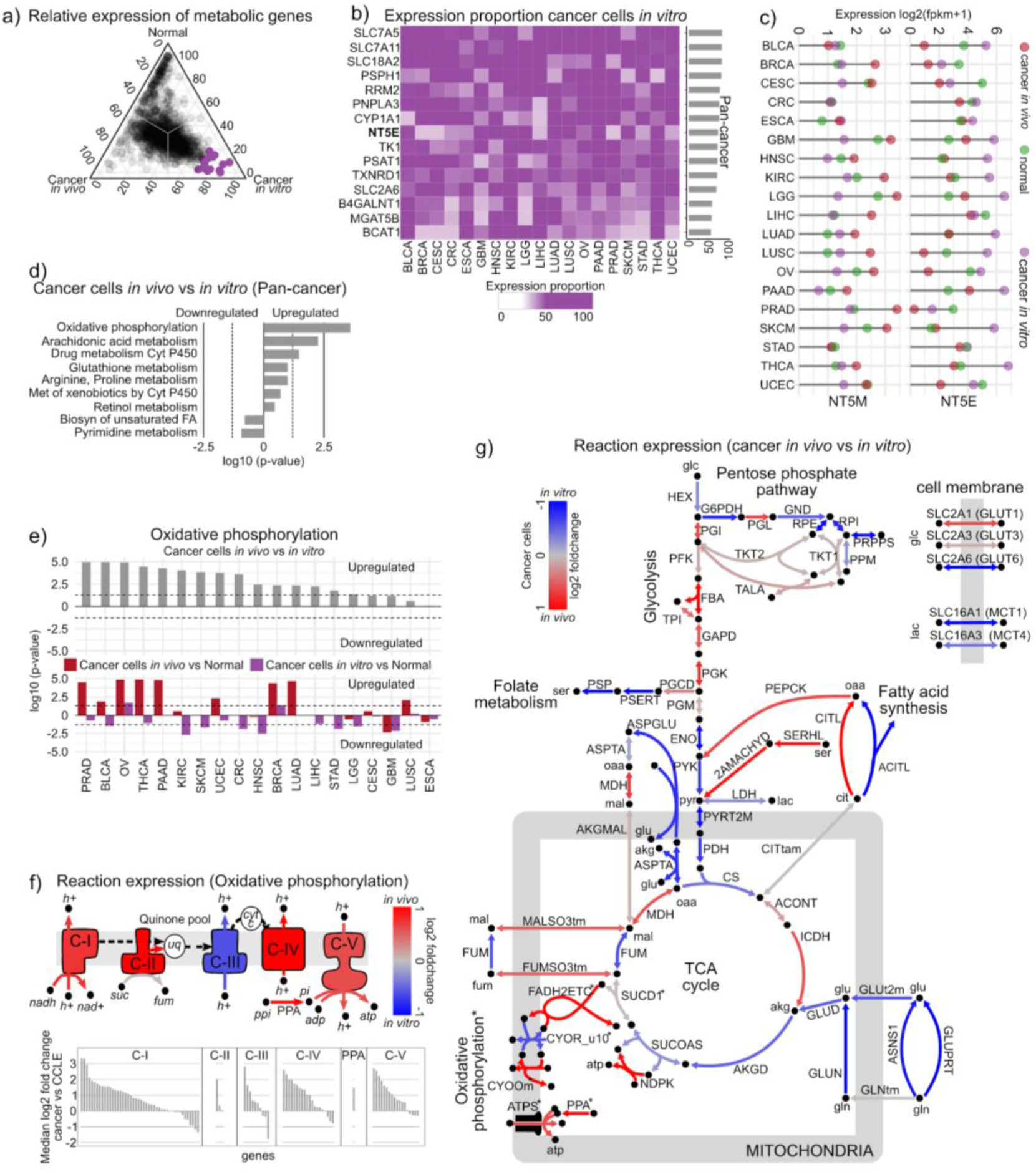
Differential metabolic pathway activity of cancer cells in vivo versus in vitro: **a)** Relative expression (proportion) of metabolic genes in cancer cells *in vivo*, *in vitro* and corresponding normal tissue across tumor types. The top-15 overexpressed cancer cell line genes are highlighted in purple color. **b)** Relative expression profiles of the top-15 overexpressed genes in cancer cell lines as compared to cancer cells *in vivo* and normal tissue expression. **b**) Expression of *NT5E* and *NT5M* in cancer cells in vivo, cancer cell lines and normal cells across tumor types. **c)** Gene sets enriched in cancer cells in vivo vs in vitro across tumor types. **d)** Enrichment of the KEGG oxphos pathway in cancer cells in vivo as compared to cancer cells in vitro across individual tumor types. **e)** Differential expression across oxphos reactions for cancer cells in vivo vs in vitro. **f)** Metabolic map of central carbon metabolism for cancer cells *in vivo* vs *in vitro*.

Analysis of genes which were differentially expressed in cancer cells *in vivo* as compared to the cancer cell lines revealed a >4 fold upregulation of *NT5E* (ecto-5’-nucleotidase) in cancer cell lines across 11 out of 20 tumor types (Fig. 5a). However, the mitochondrial isoform of 5’-nucleotidase (*NT5M*) shows >2 fold upregulation in cancer cells *in vivo* as compared to cancer cell lines in 10 out of 20 tumor types (Fig. 2b and 5b). Ecto 5’ nucleotidase converts extracellular nucleoside monophosphate into bioactive nucleoside intermediates and has been linked to absorption of deoxyribonucleotide, especially dAMP (Narumi et al. 2019). However, the mitochondrial 5’-nucleotidase plays an important role in maintaining dTTP levels in mitochondria and thus preventing malfunction of mitochondrial DNA synthesis (Gallinaro et al. 2002). This observation reflects that while cancer cells *in vitro* utilize ecto-5’-nucleotidase to utilize extracellular metabolites in their media to support growth, cancer cells *in vivo* use the mitochondrial isoform of this enzyme to protect mitochondrial DNA replication.

Next, we conducted a pan-cancer analysis of KEGG metabolic pathways, contrasting *in vivo* and *in vitro* cancer cell gene expression levels (See Methods). Interestingly, this analysis highlighted higher *in vivo* activity of the CYP450 pathway (Fig. 5c), involved in metabolism and turnover of drugs and hormones (Fatunde and Brown 2020), potentially reflecting the more heterogeneous and challenging environment of cancer cells *in vivo*. Strikingly, we also observed significant upregulation of Oxphos in cancer cells *in vivo* as compared to *in vitro* (Fig. 5c). Oxphos upregulation *in vivo* was found across almost all tumor types (Fig. 5d & Suppl. Fig. 7). To explore reaction-level differences, we estimated and compared RECON3 reaction capacities for cancer cells *in vitro* and *in vivo* for all tumor types except brain tumors (LCG and GBM, consistent with the previous *in vivo* analysis). This reaction-level analysis further supported increased capacity of 4 out of 5 Oxphos components *in vivo* (Fig. 5e). A more detailed inspection of central carbon metabolism also showed increased expression of pyruvate kinase and lactate dehydrogenase *in vitro* (Fig. 5f). Overall, this data suggest that cultured cancer cells may rely more on glycolysis for energy production, even in the presence of oxygen, supporting a general paradigm where cancer cells *in vitro* display a more Warburg-like phenotype than cancer cells *in vivo*.

## Discussion

Profiling of bulk tumor gene expression is commonly used to study cancer metabolism. Here, using a bulk tumor transcriptome deconvolution approach, we show that metabolic gene expression signatures vary substantially between cancer and non-cancer (stromal) cells inside the tumor microenvironment. Using this data, we highlight metabolic processes that are preferential active in cancer cells relative to stromal cells and normal tissue.

Our data suggests that genes involved in dTTP synthesis, *RRM2*, *TYMS* and *TK1,* are overexpressed in cancer cells *in vivo*, likely indicating a higher demand for dTTP used for cell division. This observation is consistent with the observation that dTTP pool expansion is associated with cell division (Stover and Weiss 2012). The dTTP pool is maintained at very low levels in differentiated cells and its size has been shown to increase 20-fold during cell division, by upregulation of *RRM1/RRM2*, *TYMS* and *TK1* during cell division (Mathews 2006; Ke et al. 2005). Interestingly, control of the dTTP pool size has been shown to be essential for genomic stability (Ke et al. 2005), suggesting that an elevated dTTP pool may additionally support tumorigenesis through increased genomic instability. *NT5M*, a mitochondrial nucleotidase recurrently upregulated in cancer cells across tumor types, may counteract this imbalance and protect the mitochondrial genome from dTTP mediated genomic instability. Dihydroorotate dehydrogenase (*DHODH*), thymidylate synthase (*TYMS*) and dihydrofolate reductase (*DHFR*) are three enzymes in pyrimidine synthesis that are commonly targeted by anticancer drugs (Christensen et al. 2019; Luengo, Gui, and Vander Heiden 2017).

The mitochondrial dicarboxylate carrier, *SLC25A10*, was markedly upregulated in cancer cells across most tumor types. *SLC25A10* transports malate and succinate through the mitochondrial membrane, regulating NADPH production and cellular metabolism. *SLC25A10* has previously been implicated in transport of mitochondrial glutathione (mGSH) to maintain antioxidant defense in cancer cells (Kamga, Zhang, and Wang 2010; Zhou et al. 2015). Strikingly, *SLC25A10*-inhibition has been shown to improve treatment outcome in combination with radiation therapy, especially in solid tumors with hypoxic cell fractions (Hlouschek et al. 2018).

We observed that the reported overexpression of *IDO1* in tumors derive from stromal rather than cancer cells. This finding is consistent with the observation that dendritic cells and macrophages express *IDO1*, which converts the amino acid tryptophan to kynurenine and helps in establishing immunologic tolerance (Munn and Mellor 2016). There is also evidence supporting that mesenchymal stromal cells within tumor microenvironment express *IDO1* (Günther, Däbritz, and Wirthgen 2019). A number of clinical trials are targeting *IDO1* for cancer treatments (Zhai et al. 2018). Our data suggests that it is the stromal cells, and not the cancer cells, within the tumor microenvironment that these therapies are modulating. Additionally, tryptophan catabolism that is shown by our data to be upregulated in stromal cells, is emerging as a target in cancer immunotherapy (Opitz et al. 2020).

A common perception is that cancer cells, even in the presence of oxygen, rely on glycolysis for energy generation and have reduced mitochondrial oxidation, the “Warburg effect” (Warburg 1956; Gandhi and Das 2019). However, recent studies indicate that mitochondrial respiration and glycolysis are not mutually exclusive and mitochondrial respiration is in many cases required for tumor growth (DeBerardinis and Chandel 2016; Xu et al. 2015). Here, we observe that oxidative phosphorylation is upregulated in cancer cells as compared to stromal cells, whereas glycolytic activity in the two is comparable. Total glucose uptake capacity of stromal cells is also higher than that of cancer cells. These results are in line with recent studies that indicate mitochondrial respiration and glycolysis are not mutually exclusive and in some cases, mitochondrial respiration may be required for tumor growth (DeBerardinis and Chandel 2016). Our data do not oppose that tumors take up more glucose as compared to normal tissue. However, our results suggest that non-malignant (stromal) cells in the tumor microenvironment may contribute significantly towards glucose consumption and lactate production. This observation is in line with a new paradigm in tumor metabolism, the reverse warburg effect (Fu et al. 2017). Additionally, recent studies have shown that, just like highly proliferative normal cells, cancer cells have highly active anabolic metabolism. Cancer cells use glucose and glutamine to fuel glycolysis, TCA cycle, oxphos and PPP (Martinez-Outschoorn et al. 2017). There is increasing evidence showing that oxphos is required for tumor growth in many tumor types (DeBerardinis and Chandel 2016). One study showed that both glycolysis, oxphos and the TCA cycle are enhanced in non-small cell lung tumors relative to adjacent benign lung tissue (Hensley et al. 2016). Cancer cells in tumors have been previously shown to take up lactate from the surroundings and convert it to pyruvate to feed TCA cycle (Martinez-Outschoorn et al. 2017).

Finally, through large-scale comparison of cancer cell metabolic profiles *in vitro* and *in vivo*, we show that oxphos is generally down regulated *in vitro*. This observation highlights the metabolic differences between cancer cells *in vitro* and *in vivo* owing to supraphysiological concentrations of metabolites like glucose, glutamine or pyruvate in culture media (Ackermann and Tardito 2019).

Using a systematic data-driven approach, we depict the metabolic phenotypes of cancer and stromal cells inside human tumors. Despite that cancer and stromal cells coexist inside the same tumor microenvironment and have access to the same resources, our study highlights key differences in their metabolism. We present a pan-cancer resource that provides new insights into tumor metabolism and enables the discovery of metabolic dependencies of cancer cells *in vivo*. By jointly inferring the metabolic profiles of cancer and stromal cells across tumor types, our results could also provide a foundation for studies of metabolic crosstalk between malignant and non-malignant cells in the tumor microenvironment.

## Methods

### Data Collection and Normalisation

Gene expression data for all TCGA, GTEX and CCLE samples was downloaded from the UCSC Xena platform (Goldman et al. 2019). For normal tissue expression, adjacent normal samples from TCGA were used for 18 tissue types. Normal brain tissue samples from the GBM study were used for LGG as well. For ovarian cancer (OV), normal ovarian samples were obtained from GTEX. TCGA tumor copy number variation data was also downloaded from Xena. Tumor purity (cancer cell fraction) estimates for all TCGA samples were obtained from a previous study (Ghoshdastider et al. 2019), where purity was jointly estimated using mutation allele frequencies, copy number profiles, and RNA signatures of each tumor sample.

Genes with zero expression in >90% of the samples were excluded from the analysis. For genes with multiple transcripts, expression of each transcript was summed to obtain gene expression. Upper quartile normalization was performed to make the expression profiles comparable across tumor types and cohorts. A genome scale metabolic model, RECON3 (Brunk et al. 2018, 3), was used to filter metabolic genes for further analysis. For all subsequent analysis, gene expression data was log transformed, log2(fpkm+1), unless specified.

### Tumor transcriptome deconvolution of cancer and stroma expression

In order to estimate the cancer and stromal cell gene expression across tumor types, we performed tumor transcriptome deconvolution as previously reported (Ghoshdastider et al. 2019). Briefly, tumor purity was estimated using genomic and RNA data available for each TCGA tumor sample and cancer and stromal compartment expression was estimated using non-negative least squares regression. We deconvoluted the transcriptomes in 20 solid tumor types comprising 7865 TCGA samples; deconvolution analysis was performed on each tumor type separately to obtain cancer and stroma expression estimates for each tumor type.

### Gene expression analysis

In order to study differentially expressed genes across cancer, stroma and normal tissue, we computed the relative expression for each cell type (cancer, stroma and normal tissue): the expression proportion for a given gene and cell type *c* was defined as the expression in *c* divided by the total expression across all three cell types. Expression proportions were computed separately for each metabolic gene and tumor type. To investigate patterns shared across tumor types (pan-cancer analysis), we calculated the proportions of median expression in each cell type for a given gene across tumor types. Gene set enrichment was performed on 186 KEGG pathways using the R Bioconductor package “fgsea” (Sergushichev 2016). In order to study differentially expressed metabolic pathways, we filtered metabolic pathways using KEGG pathway IDs.

### Metabolic map generation

To study metabolic functions of cancer and stromal cells at the reaction level, we integrated deconvoluted cancer and stroma expression with a human metabolic model, RECON3, the most comprehensive genome-scale model containing metabolite and protein information of metabolic functions in humans (Brunk et al. 2018). Additionally, RECON3 contains information of genes and proteins associated with each metabolic reaction in the form of boolean rules. These Gene-Protein-Reaction (GPR) associations describe subunits of a protein complex using “AND” operators and isozymes using “OR” operators. GPRs are the access points for incorporation of genetic, transcriptomic or proteomics data into metabolic networks (Swainston et al. 2016; Richelle, Joshi, and Lewis 2018). Ensembl gene IDs in RNA-seq dataset were mapped to Entrez IDs used in RECON3 using R Bioconductor packages, biomaRt and org.Hs.eg.db (Durinck et al. 2009; Carlson 2019). The deconvoluted cancer and stromal cell expression was integrated with RECON3 based on the GPR information. To estimate reaction expression from the gene expression data, we evaluated the GPR rule of each reaction while replacing the “OR” and “AND” operators with “max” and “min”, respectively (Auslander et al. 2016; Richelle, Joshi, and Lewis 2018). Although RECON3 is one of the most comprehensive and curated human metabolic models, there are still some gaps in the model. Thus, manual verification and curation is an important step in model generation (Duarte et al. 2007). While looking at oxidative phosphorylation we found that the components of two reactions, CYOOm3i (complex IV) and ATPS4mi (complex V), had incorrect isoform annotation. A literature search was used to manually curate these reactions (see Supplementary Methods). Escher maps were used to visualise the metabolic networks (King et al. 2015).

## Data Availability

All datasets generated in this study are available at https://camp.genome.sg/ui.

