## supplementary material for "A pan-cancer metabolic atlas of the tumor microenvironment"

**Rohatgi, et al., 2020**

**Genome Institute of Singapore, Singapore 138672, Singapore**

### Supplementary table

#### Supplementary Table 1:

| | |
|---|---|
| BLCA | Bladder Urothelial Carcinoma |
| BRCA | Breast invasive carcinoma |
| CESC | Cervical squamous cell carcinoma and endocervical adenocarcinoma |
| CRC | colorectal carcinoma |
| ESCA | Esophageal carcinoma |
| GBM | Glioblastoma multiforme |
| HNSC | Head and Neck squamous cell carcinoma |
| KIRC | Kidney renal clear cell carcinoma |
| KIRP | Kidney renal papillary cell carcinoma |
| LGG | Brain Lower Grade Glioma |
| LIHC | Liver hepatocellular carcinoma |
| LUAD | Lung adenocarcinoma |
| LUSC | Lung squamous cell carcinoma |
| OV | Ovarian serous cystadenocarcinoma |
| PAAD | Pancreatic adenocarcinoma |
| PRAD | Prostate adenocarcinoma |
| SKCM | Skin Cutaneous Melanoma |
| STAD | Stomach adenocarcinoma |
| THCA | Thyroid carcinoma |
| UCEC | Uterine Corpus Endometrial Carcinoma |

#### Supplementary Table 2:

| ID | Description | GeneRatio | BgRatio | p.adjust | geneID |
|---|---|---|---|---|---|
| GO:0006259 | DNA metabolic process | 5/27 | 32/1855 | 0.044537 | CBS/NT5M/RRM2/TK1/TYMS |
| GO:0009162 | deoxyribonucleoside monophosphate metabolic process | 3/27 | 10/1855 | 0.100772 | NT5M/TK1/TYMS |
| GO:0009262 | deoxyribonucleotide metabolic process | 3/27 | 18/1855 | 0.422581 | NT5M/RRM2/TYMS |
| GO:0090304 | nucleic acid metabolic process | 6/27 | 122/1855 | 0.574255 | CA9/CBS/NT5M/RRM2/TK1/TYMS |
| GO:0006260 | DNA replication | 2/27 | 10/1855 | 0.574255 | NT5M/RRM2 |

### Supplementary Methods

#### Manual Curation of reaction GPRs:

NADH2_u10mi (Complex I):

Reaction “NADH2_u10mi” represents mitochondrial complex I (NADH Dehydrogenase) in RECON3 and is associated with 46 genes in the model. A study published in 2012 showed that NDUFA4 that was formerly considered to be a component of complex I, is a constituent of the complex IV instead (Balsa et al. 2012). Therefore, we moved the NDUFA4 gene from complex I to complex IV. Additionally, we could not find evidence for TUSC3 to be associated with complex I and thus removed it from the GPR for this reaction. The new GPR for this reaction is as follows:

(MT-ND1 & MT-ND2 & MT-ND3 & MT-ND4 & MT-ND4L & MT-ND5 & MT-ND6 & NDUFA1 & NDUFA2 & NDUFA3 & NDUFA5 & NDUFA6 & NDUFA7 & NDUFA8 & NDUFA9 & NDUFA10 & NDUFAB1 & NDUFB1 & NDUFB2 & NDUFB3 & NDUFB4 & NDUFB5 & NDUFB6 & NDUFB7 & NDUFB8 & NDUFB9 & NDUFB10 & NDUFC1 & NDUFC2 & NDUFS1 & NDUFS2 & NDUFS3 & NDUFV1 & NDUFS4 & NDUFS5 & NDUFS6 & NDUFS8 & NDUFV2 & NDUFV3 & NDUFA13 & NDUFB11 & NDUFA12 & NDUFA11 & NDUFS7)

CYOOm3i (Complex IV):

Reaction “CYOOm3i” represents mitochondrial complex IV (COX) in RECON3. This reaction is associated with 20 genes connected to each other with the “AND” operator, which implies all these genes produce subunits essential for the formation of COX protein complex. Whereas, literature says that COX consists of 14 subunits, 3 encoded by the mitochondrial DNA and 11 by nuclear DNA (Sinkler et al. 2017). Tissue and condition specific isoforms have been reported for COX. COX4I1, COX4I2, COX6A1, COX6A2, COX6B1, COX6B2, COX7A1, COX7A2, COX7A2L, COX8A, and COX8C are tissue-specific isoforms expressed in liver, heart and skeletal muscle, lung, and testes (Sinkler et al. 2017; Hüttemann, Schmidt, and Grossman 2003). Because these isoforms perform the same function in different tissues or conditions these should be associated with each other with operator “OR”. Additionally, COX7B2 protein is inferred from homology therefore it could be an isoform of COX7B (The UniProt Consortium 2018). NDUFA4, which has been added to this reaction from complex I (see above). This information was used to manually correct the gene rule for complex IV to be:

(COX4I1 or COX4I2) and COX5A and COX5B and (COX6A1 or COX6A2) and (COX6B1 or COX6B2) and COX6C and (COX7A1 or COX7A2 or COX7A2L) and (COX7B or

COX7B2) and COX7C and (COX8A or COX8C) and NDUFA4 and COX1 and COX2 and COX3

ATPS4mi (ATP synthase):

Reaction “ATPS4mi” represents mitochondrial complex V (ATP synthase) in RECON3. This reaction contains a pseudogene ATP5MGL (ATP Synthase Membrane Subunit G Like) connected with “AND” to other genes, which means that this protein is a necessary part of the protein complex. In contrast, according to the Uniprot database the product of this gene is annotated due to its similarity with another subunit of ATP synthase, ATP5MG (The UniProt Consortium 2018). Therefore, we have modified the GPR in a way that ATP5MGL “OR” ATP5MG are able to produce subunit of the protein complex.

### Supplementary Figures

#### Supplementary Fig 1: Transcriptome deconvolution of genes overexpressed in cancer cells

Gene expression data for dTTP pathway specific genes (*RRM2*, *TK1*, *TYMS*) as well as the mitochondrial factors *SLC25A10* and *NT5M*. Gene expression (y-axis, log2) and tumor purity (x-axis) is displayed for each gene across 20 solid tumor types. Each data point is a single tumor sample. Blue and red dots at the ends of regression lines show the inferred stromal and cancer expression for the given gene and tumor type. Data points are colored according to the DNA copy number level (deletion vs. amplification) at the given gene locus.

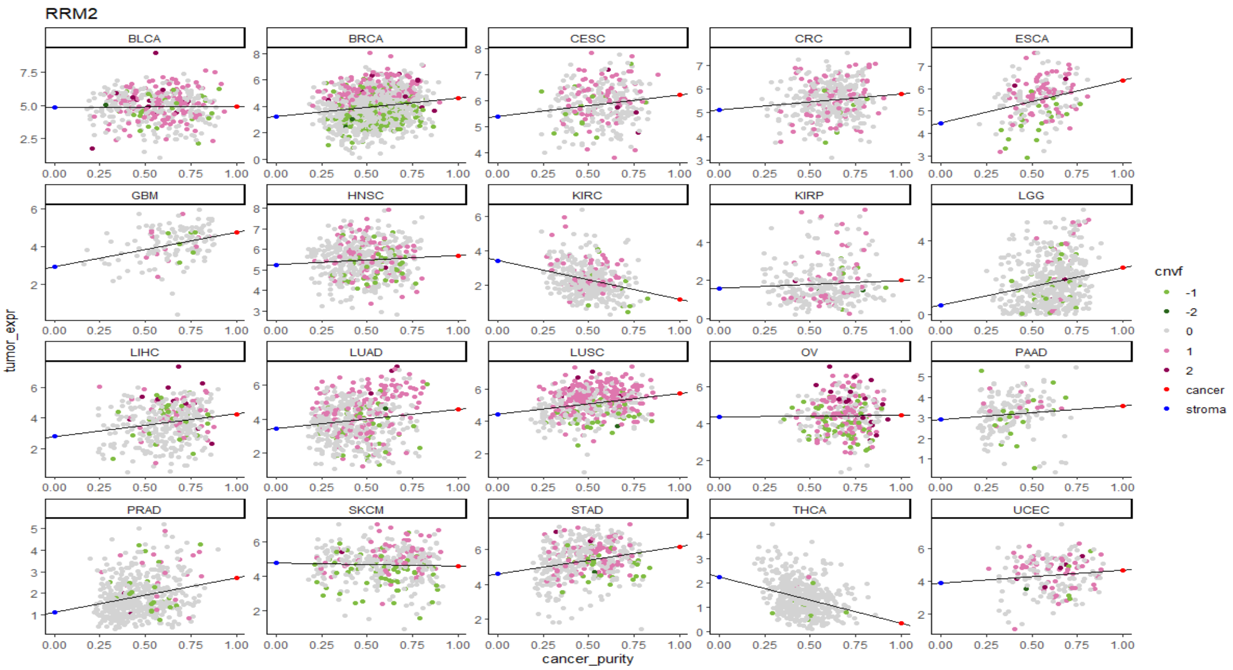

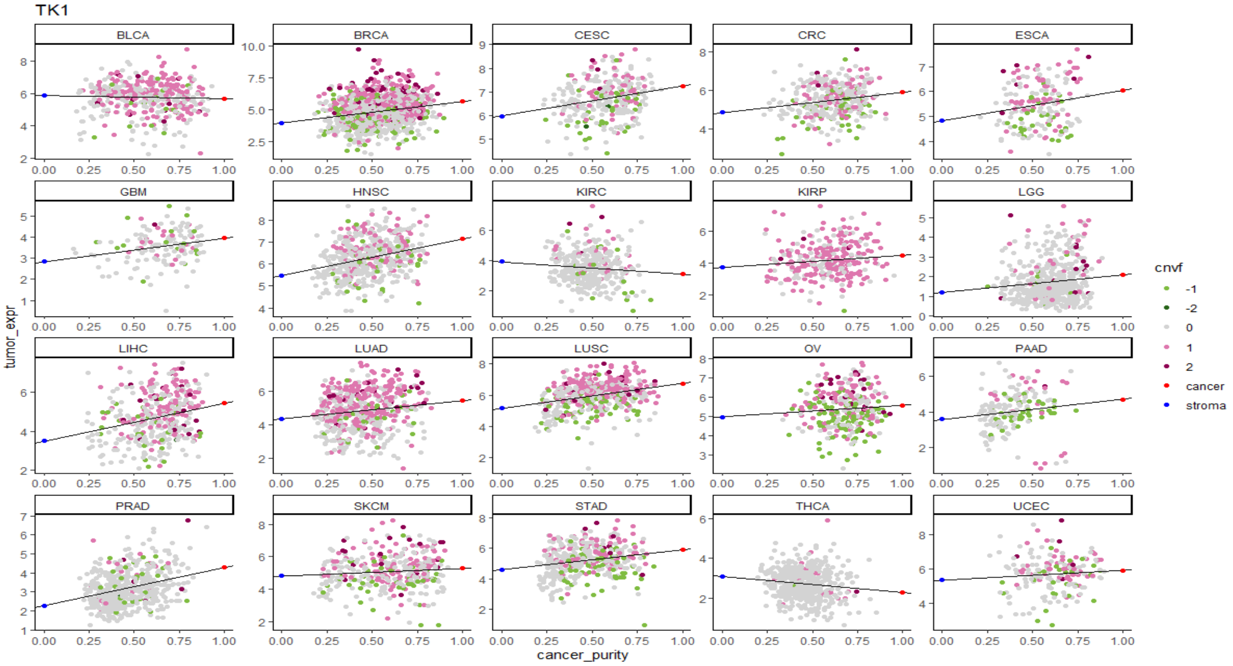
TK1
BLCA
BRCA
CESC
CRC
ESCA
GBM
HNSC
KIRC
KIRP
LGG
LIHC
LUAD
LUSC
OV
PAAD
PRAD
SKCM
STAD
THCA
UCEC
tumor_expr
cancer_purity
cnvf
-1
-2
0
1
2
cancer
stroma

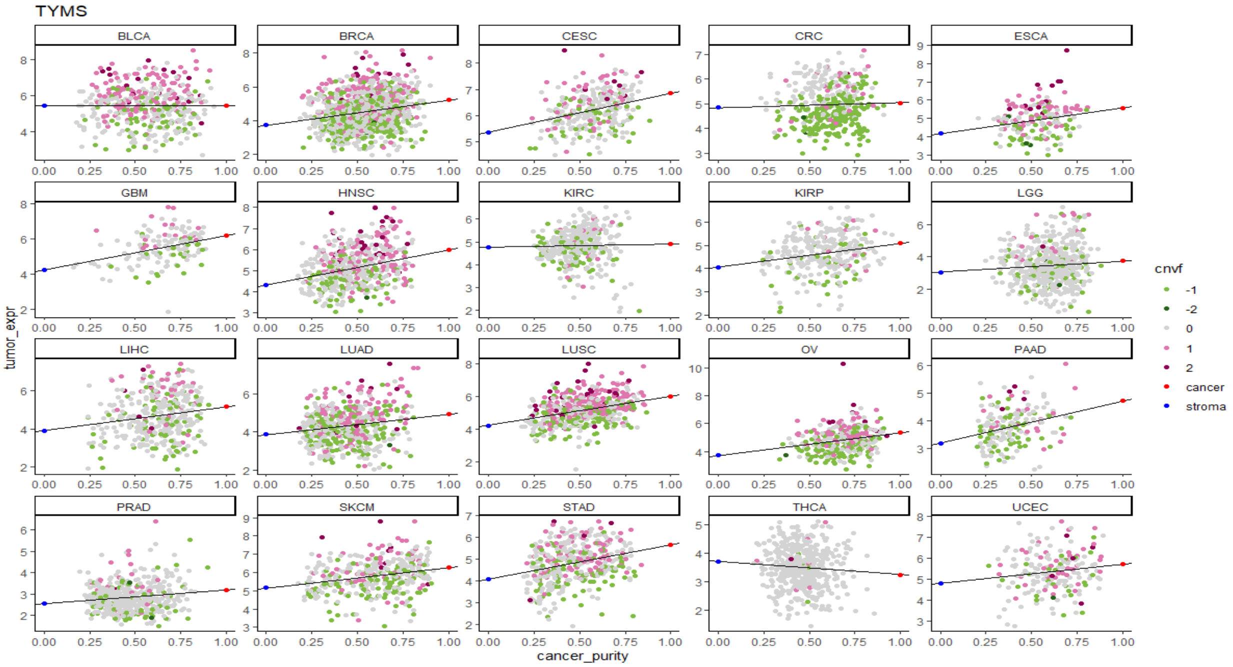
TYMS
BLCA
BRCA
CESC
CRC
ESCA
GBM
HNSC
KIRC
KIRP
LGG
LIHC
LUAD
LUSC
OV
PAAD
PRAD
SKCM
STAD
THCA
UCEC
tumor_expr
cancer_purity
cnvf
-1
-2
0
1
2
cancer
stroma

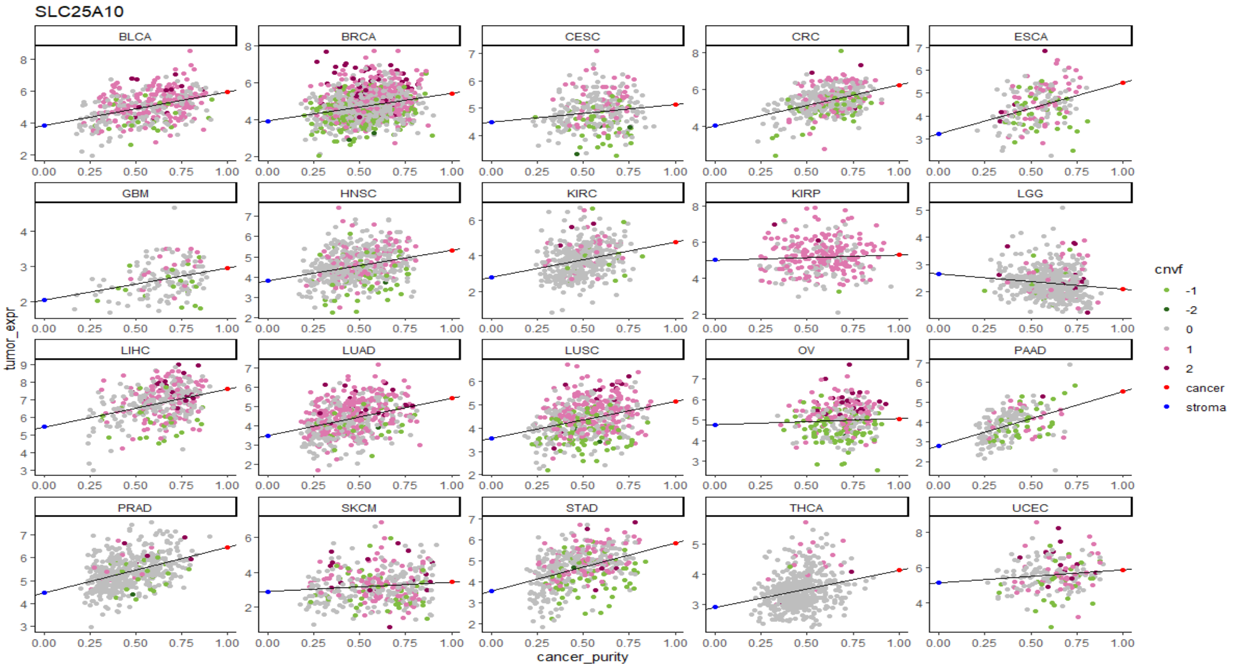
SLC25A10
BLCA
BRCA
CESC
CRC
ESCA
GBM
HNSC
KIRC
KIRP
LGG
LIHC
LUAD
LUSC
OV
PAAD
PRAD
SKCM
STAD
THCA
UCEC
tumor_expr
cancer_purity
cnvf
-1
-2
0
1
2
cancer
stroma

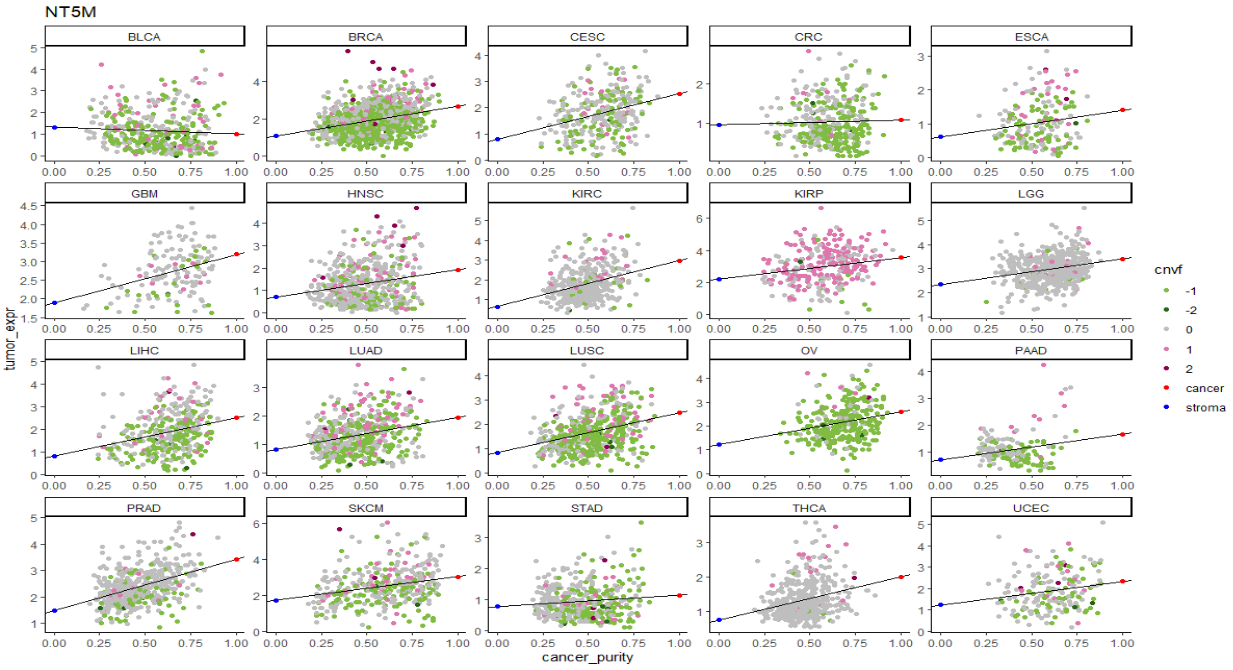
NT5M
BLCA
BRCA
CESC
CRC
ESCA
GBM
HNSC
KIRC
KIRP
LGG
LIHC
LUAD
LUSC
OV
PAAD
PRAD
SKCM
STAD
THCA
UCEC
tumor_expr
cancer_purity
cnvf
-1
-2
0
1
2
cancer
stroma

#### Supplementary Fig 2: Expression of cancer-cell specific genes

Bulk tumor expression of genes with inferred overexpression in cancer cells (*RRM2*, *TK1*, *TYMS*, *NT5M* and *SLC25A10*) across different tumor types, compared to corresponding normal tissues. The cancer and stroma data points are inferred using tumor transcriptome deconvolution.

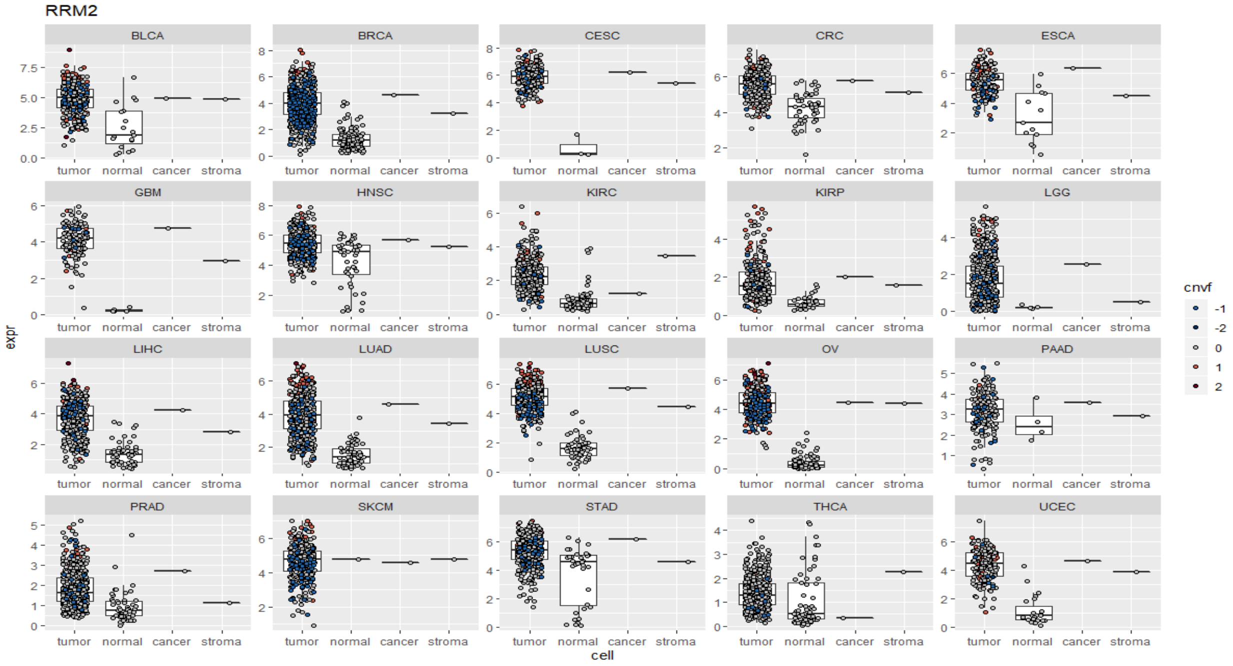

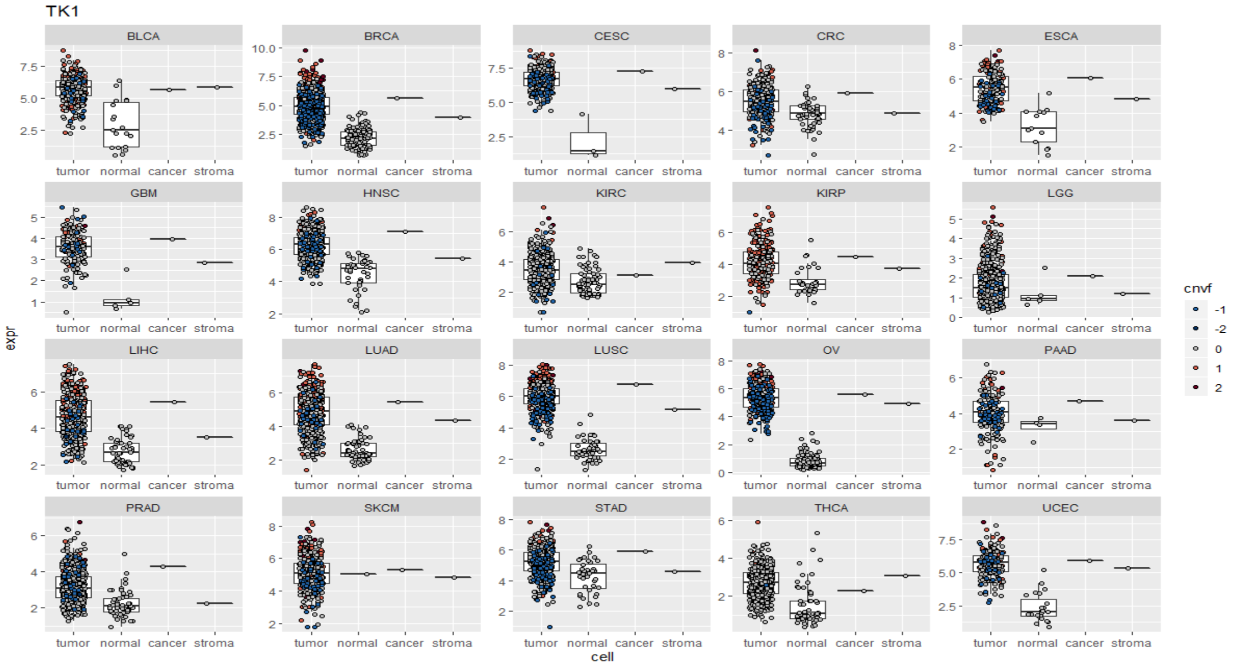

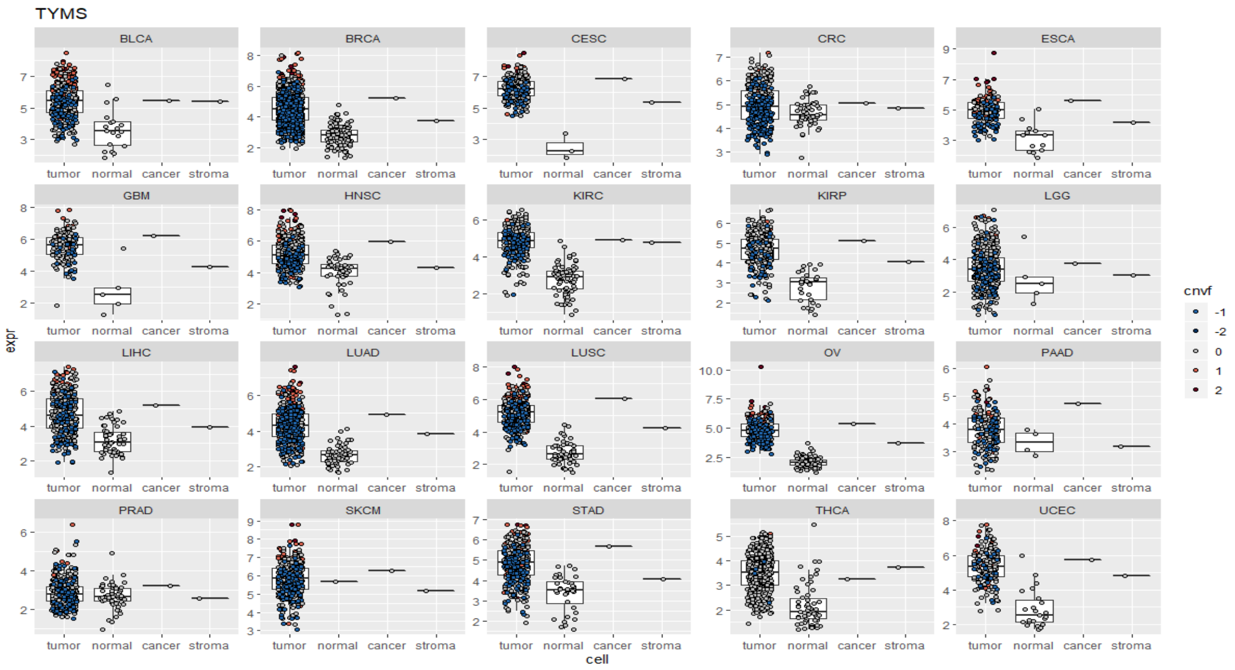
TYMS
BLCA
BRCA
CESC
CRC
ESCA
GBM
HNSC
KIRC
KIRP
LGG
LIHC
LUAD
LUSC
OV
PAAD
PRAD
SKCM
STAD
THCA
UCEC
tumor normal cancer stroma
expr
cell
cnvf
-1
-2
0
1
2

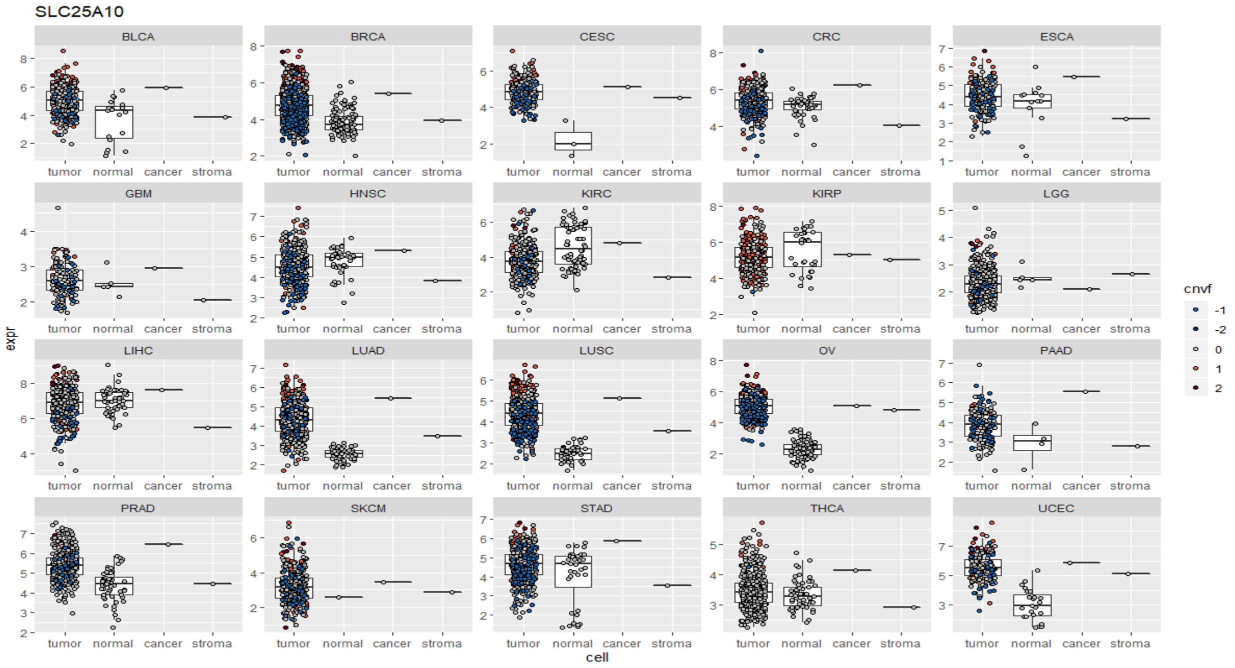
SLC25A10
BLCA
BRCA
CESC
CRC
ESCA
GBM
HNSC
KIRC
KIRP
LGG
LIHC
LUAD
LUSC
OV
PAAD
PRAD
SKCM
STAD
THCA
UCEC
tumor normal cancer stroma
expr
cell
cnvf
-1
-2
0
1
2

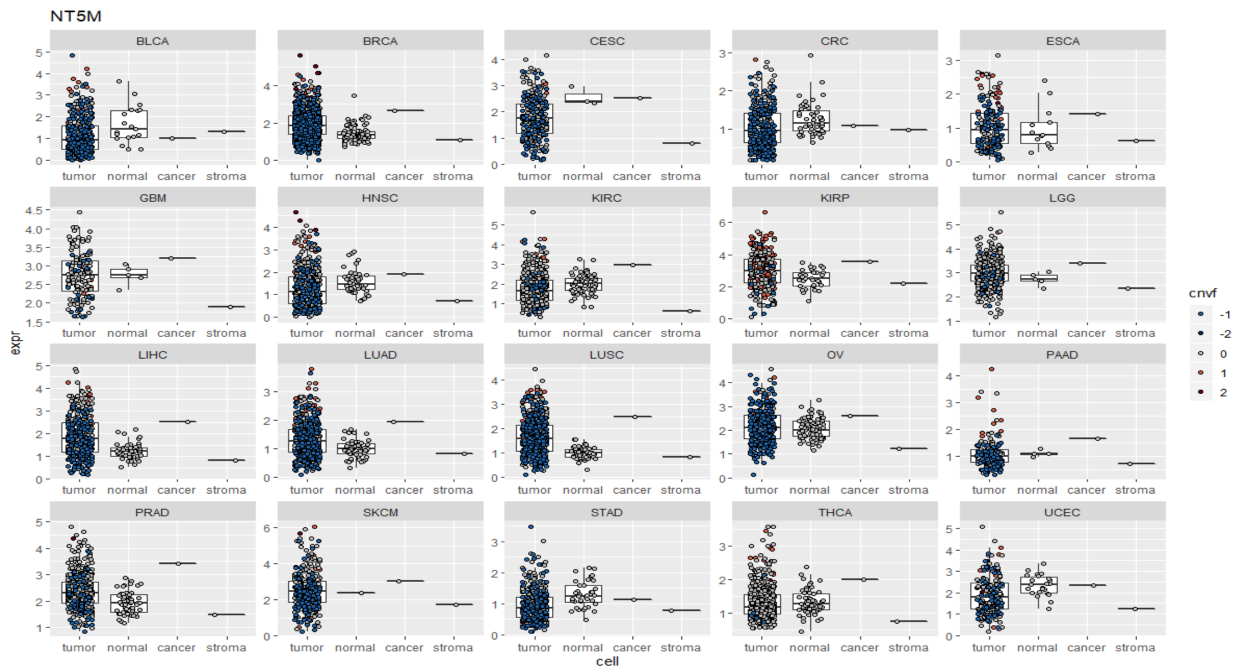

#### Supplementary Fig 3: Transcriptome deconvolution of *IDO1*

Gene expression data for *IDO1*. Gene expression (y-axis, log2) and tumor purity (x-axis) is displayed for each gene across 20 solid tumor types. Each data point is a single tumor sample. Blue and red dots at the ends of regression lines show the inferred stromal and cancer expression for the given gene and tumor type. Data points are colored according to the DNA copy number level (deletion vs. amplification) at the given gene locus.

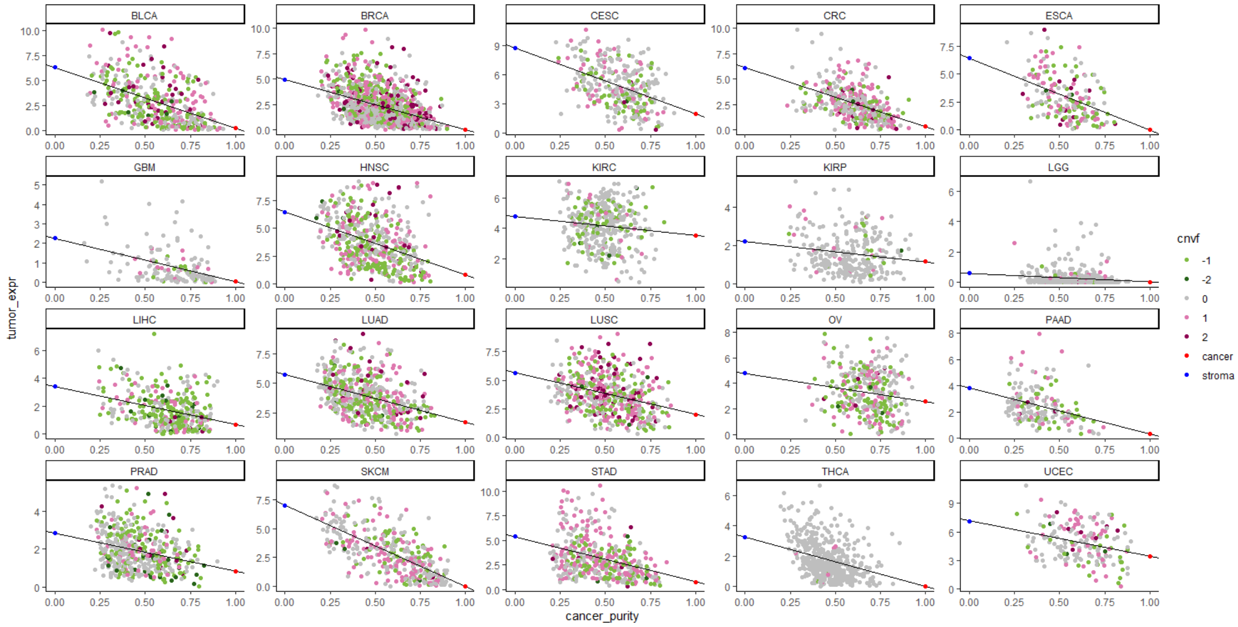

#### Supplementary Fig 4: IDO1 expression in stroma

Bulk tumor expression of stroma-specific *IDO1* across different tumor types, compared to corresponding normal tissues. The cancer and stroma data points are inferred using tumor transcriptome deconvolution.

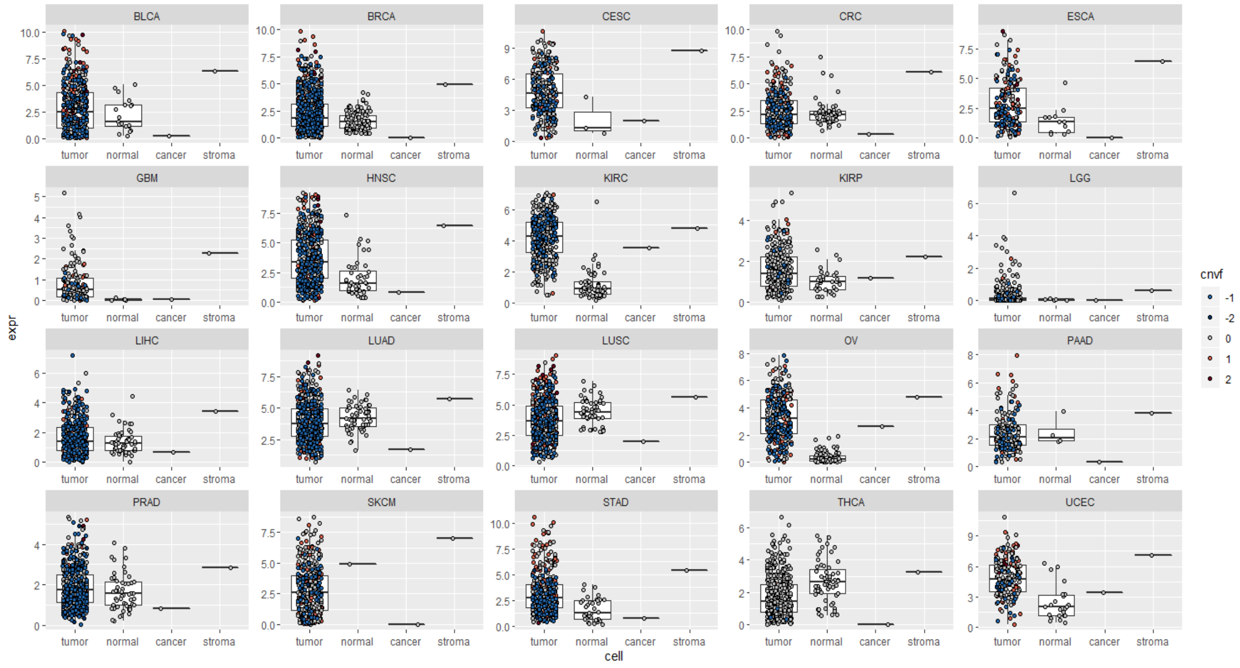

#### Supplementary Fig 5: Oxphos GSEA, cancer vs. stroma

Running enrichment score (ES) of KEGG oxidative phosphorylation for cancer vs. stroma expression across tumor types, obtained with GSEA.

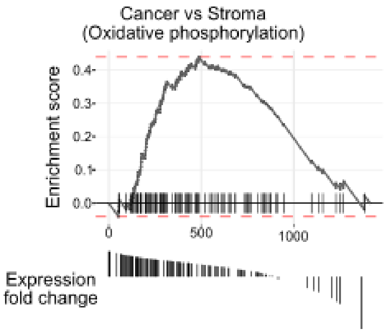

**Supplementary Fig 6: RECON3 metabolic map for brain cancer (Cancer vs Stroma)**

Metabolic maps of central carbon metabolism. For glioblastoma (GBM) and lower grade glioma (LGG), cancer (left) and stromal (right) expression estimated for each reaction was compared to normal cells.

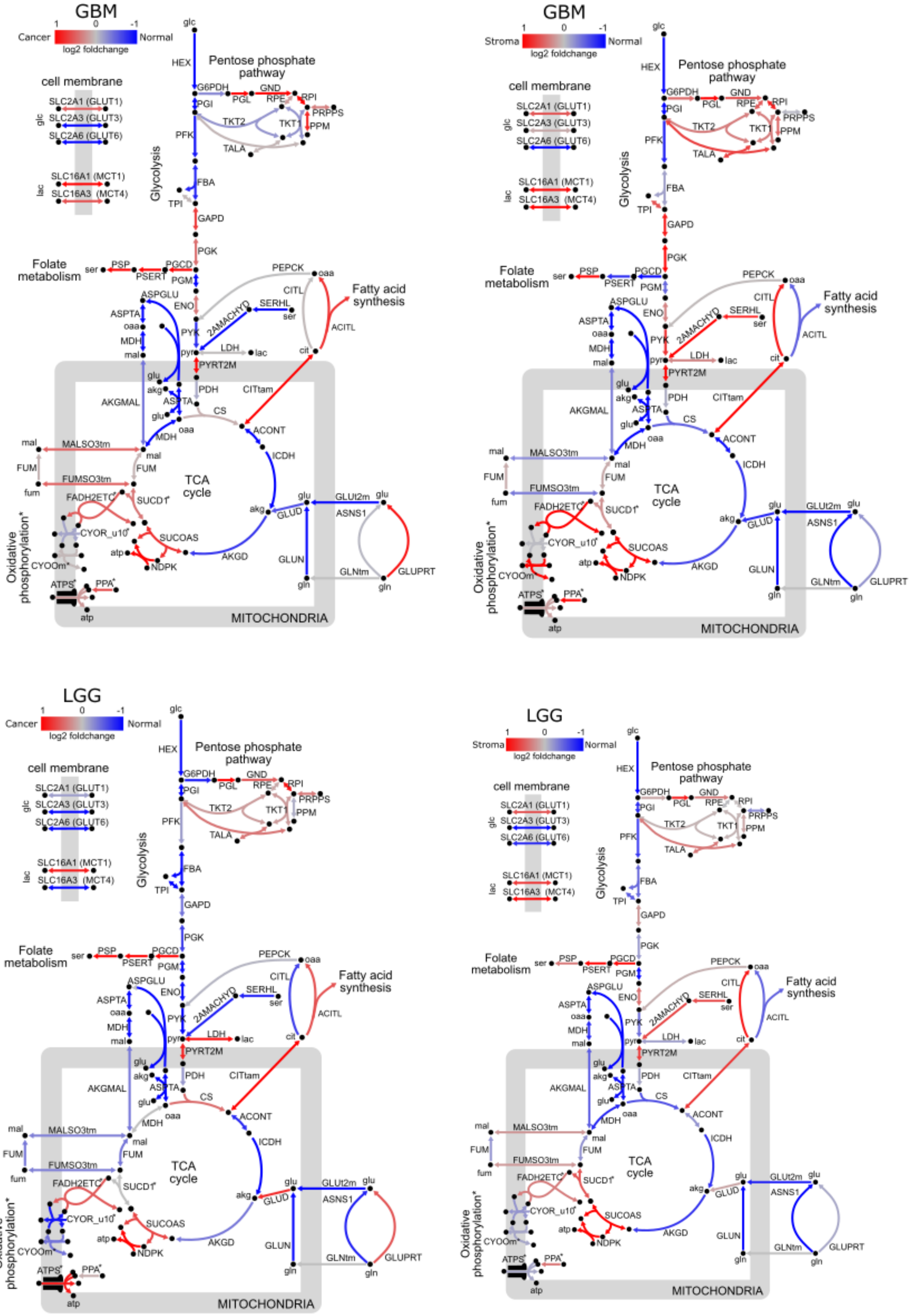

**Supplementary Fig 7: GSEA of Oxphos, Cancer *in vivo* vs *in vitro***

Running enrichment score (ES) of KEGG oxidative phosphorylation for cancer *in vivo* vs *in vitro*, obtained by GSEA

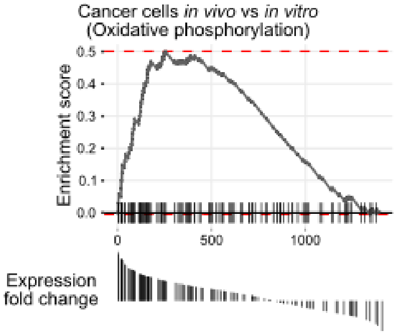
